# Potent, Gut-Restricted Inhibitors of Divalent Metal Transporter 1 (DMT1): Preclinical Efficacy Against Iron Overload and Safety Evaluation

**DOI:** 10.1101/2022.09.01.506269

**Authors:** Alison Cutts, Sultan Chowdhury, Laszlo G Ratkay, Maryanne Eyers, Clint Young, Rostam Namdari, Jay A Cadieux, Navjot Chahal, Michael Grimwood, Zaihui Zhang, Sophia Lin, Ian Tietjen, Zhiwei Xie, Lee Robinette, Luis Sojo, Matthew Waldbrook, Michael Hayden, Tarek Mansour, Simon Pimstone, Y. Paul Goldberg, Michael Webb, Charles J. Cohen

## Abstract

Divalent metal transporter 1 (DMT1) cotransports ferrous iron and protons and is the primary mechanism for uptake of non-heme iron by enterocytes. Inhibitors are potentially useful as therapeutic agents to treat iron overload disorders such as hereditary hemochromatosis or β-thalassemia intermedia, provided that inhibition can be restricted to the duodenum. We used a calcein quench assay to identify human DMT1 (hDMT1) inhibitors. Dimeric constructs were made to generate more potent compounds with low systemic exposure. Direct block of DMT1 was confirmed by voltage clamp measurements. The lead compound, XEN602, strongly inhibits dietary iron uptake in both rats and pigs, yet has negligible systemic exposure. Efficacy is maintained for >2 weeks in a rat subchronic dosing assay. Doses that lowered iron content in the spleen and liver by >50% had no effect on the tissue content of other divalent cations except for cobalt. XEN602 represents a powerful pharmacological tool for understanding the physiological function of DMT1.

**SIGNIFICANCE STATEMENT:** This report introduces methodology to develop potent, gut-restricted inhibitors of DMT1 and identifies XEN602 as a suitable compound for in vivo studies. We also report novel animal models to quantify the inhibition of dietary uptake of iron in both rodents and pigs. This research shows that inhibition of DMT1 is a promising means to treat iron overload disorders.

## INTRODUCTION

Iron is an essential nutrient, but excess iron is toxic because it catalyzes reactions which release damaging free radicals. Mammals have no physiological excretory mechanisms for the elimination of excess iron and therefore rely on very tightly regulated uptake from dietary sources. Several genetic disorders are known, in both laboratory animals and humans, that identify divalent metal transporter 1 (DMT1, also known as natural resistance-associated macrophage protein 2 [NRAMP2], divalent cation transporter 1 (DCT1) and solute carrier family 11, member 2 [SLC11A2]) as the primary transport mechanism for uptake of non-heme iron by enterocytes (Andrews and Schmidt, 2007). DMT1 is a symporter for protons and ferrous iron (Gunshin et al., 1997). Excessive iron uptake is normally prevented by regulation of DMT1 expression. Inappropriate overexpression of this protein may contribute to the iron overload burden in multiple disorders, most significantly hereditary hemochromatosis (HH) and β-thalassemia intermedia (Fleming and Ponka, 2012). Existing treatments for these disorders do not directly inhibit DMT1 or reduce iron uptake from the gut and have significant limitations or adverse side effects.

HH is the most common genetic disorder among people of European descent (Fleming and Ponka, 2012). HH usually results from a defect in the *HFE* gene that results in excess DMT1 expression and excess non-heme iron uptake (Lynch et al., 1989; Kelleher et al., 2004). Currently, the standard treatment for HH is repeat phlebotomy. There is a medical need for a better treatment because in some patients compliance might be poor, and, importantly, the phlebotomy can trigger increased erythropoiesis and iron uptake via intestinal DMT1.

Iron overload is also an important consequence of β-thalassemia intermedia (Gardenghi et al., 2010). These patients suffer a milder form of anemia than those that have thalassemia major and thus do not require transfusions on a regular basis, even though they are anemic. Intestinal DMT1 is upregulated due to the ineffective erythropoiesis and anemia (Taher et al., 2009), resulting in intestinal iron hyperabsorption and, eventually, iron overload. Phlebotomy is not an option for patients with thalassemia (as they are anemic), and the present standard of care is the use of iron chelators such as desferrioxamine. However, these agents have potential for serious dose-limiting side effects and usually have poor oral bioavailability.

Direct blockade of DMT1-mediated iron influx in enterocytes by an orally administered small-molecule inhibitor is likely to provide an improved therapeutic approach for the treatment of iron overload. However, DMT1 is also expressed in erythroid cells, macrophages, neurons and hepatocytes, and inhibition of transport in these tissues could potentially compromise systemic iron homeostasis (Garrick and Garrick, 2009). Missense mutations in DMT1 have been characterized in microcytic anemia (*mk*) mice (Fleming et al., 1997), Belgrade rats (Fleming et al., 1998) and humans (Priwitzerova et al., 2005) and in all cases result in a microcytic anemia. Similarly, knockout of DMT1 in mice indicates that iron uptake via DMT1 in reticulocytes is essential for normal erythropoiesis (Gunshin et al., 2005). Therefore, limiting the exposure of a DMT1 inhibitor compound to the gut (i.e., minimizing systemic bioavailability) is likely essential for adequate safety.

Previously reported small-molecule inhibitors of DMT1 are orally efficacious in animal models of iron overload, but the doses needed for therapeutic effect and the systemic exposure are higher than those acceptable for therapeutic development (Brown et al., 2004; Wetli et al., 2006; Cadieux et al., 2012; Zhang et al., 2012). Here, we report the characterization of compounds developed using a strategy of covalently linking monomers to form dimers with greatly improved potency and very low systemic exposure. This strategy has been previously deployed to good effect in the development of gut-restricted potent inhibitors of the cystic fibrosis transmembrane conductance regulator (CFTR) (Sonawane et al., 2008).

In order to quantify block of DMT1 transport in vitro, we have used a combination of a fluorescence assay for divalent cation influx and voltage clamp studies of DMT1 (Breuer et al., 1995; Picard et al., 2000; Xu et al., 2004). Furthermore, we have developed both acute and subchronic in vivo assays to assess DMT1 activity in rats. Here we describe the efficacy of compounds in blocking iron uptake by the transporter in both types of assay. Iron may be absorbed from the gut by at least two mechanisms, depending on whether it is elemental or heme bound, and the relative contributions of these two processes vary as a function of diet and physiological status of the animal (Shayeghi et al., 2005). Since laboratory rats have a diet that is very low in heme-containing protein, the proportion of iron uptake due to DMT1 in rats may not be representative of that in larger, non-rodent animals that have a more varied diet. We therefore established an iron hyperabsorption model in weanling pigs and find similar potency of our lead compounds in pigs as in rats. We further our in vivo assessment of our lead compound by exploring safety and toxicity of higher doses in the rat.

## MATERIALS AND METHODS

### Procedures for the Synthesis of Compounds (4a, 4b, 5–21)

Compounds **4a, 4b** and **5–21** were synthesized utilizing procedures described previously (Zhang et al., 2012). Further details and properties are provided in the supplementary material.

### Cell Lines and Cell Culture

T-REx-CHO (Chinese hamster ovary) cells stably express the tetracycline (Tet) repressor and are designed for use with Tet-regulated expression systems according to the manufacturer’s instructions (Thermofisher). Selection of stably transfected clones was performed using 400 μg/ml of zeocin. Individual colonies were then isolated and expanded. hDMT1 expression was induced by growing the cells in medium containing 1 μg/ml of Tet for 24 h.

### In Vitro Assays

#### Calcein Quench

The calcein quench assay was performed as described previously (Cadieux et al., 2012).

#### Voltage Clamp Measurements

Whole-cell voltage clamp studies of heterologously expressed hDMT1 used a modification of a published protocol (Xu et al., 2004). From a holding potential of 0 mV, currents were elicited at 1 Hz by stepping to −150 mV for 40 ms followed by a 100 ms ramp to +60 mV, then holding at +60 mV for 40 ms before stepping back to 0 mV. As there is only charge carrier for inward current, the currents are strongly inward rectified, as reported previously (Xu et al., 2004). Consequently, the current measured while holding the voltage at −150 mV represents the influx of protons and divalent cations via DMT1 while the current measurements at +60 mV indicate a non-specific increase in leak current. Background leak currents were measured at pH 7.4 (baseline). Current due to DMT1 was taken as the difference between background and that measured at low pH with or without the addition of divalent cations. The baseline-corrected drug response was fitted to the Hill equation to generate IC_50_ values. The pipette solution contained (in mM): Cs methanesulfonate, 120; Na2ATP, 2; HEPES, 20; EGTA, 10; NaCl, 4; MgCl_2_, 2; pH was 7.2, adjusted with CsOH. The extracellular solution contained (in mM): N-methyl-D-glucamine methane sulfonate, 150; CaCl_2_, 0.2; MgCl_2_, 4; MES, 10; HEPES, 10; glucose, 10; initial pH was 7.4, adjusted with methanesulfonic acid to indicated pH.

### In Vivo Assays

#### Animals and Diet

All rat experiments were conducted on Sprague Dawley rats and were approved by the Xenon Animal Care Committee in compliance with Canadian Council on Animal Care guidelines. Normal rat diet (Formulab 5008, Harlan) contained iron at a level of 230 ppm. Where required, a rodent diet with low iron levels (2 ppm) was substituted (Harlan #TD.80396). Domestic weanling pigs were obtained and dosed by Pre-Clinical Research Services Inc. (Ft Collins, CO). Baseline levels of hemoglobin (Hb) and/or serum iron were used to determine the iron-deficient status of rats and weanling pigs. **XEN601** or **XEN602** was administered p.o. (orally) in a vehicle of 30% propylene glycol and 70% D5W (5% dextrose in H_2_O) at a volume of 4 ml/kg to both rats and weanling pigs.

#### Sample Collection and Analysis

For rat experiments involving determination of changes in serum iron levels or Hb levels, baseline blood collection was carried out 2 days before initiation of dosing after a 4 h fasting period. About 400 μl of blood was collected via jugular bleeding under Fluothane inhalation anesthesia. Similar standard procedures were used in weanling pigs to collect 2 ml samples. Blood was allowed to clot for about 1 h, and serum was prepared by centrifuging at 1300 *g* for 10 min at room temperature in an Eppendorf micro-centrifuge.

#### Serum Iron and Blood Hemoglobin Measurement

Serum iron levels were determined by a colorimetric assay. Rat samples were analyzed at IDEXX Reference Laboratories (Delta, BC, Canada) using an Olympus Analyzer. The blood Hb level for each rat was measured using a handheld HemoPoint H2 Photometer. The whole blood sample and serum sample from each weanling pig were sent to the Colorado State University Diagnostic Laboratory for complete blood count and serum iron assay, respectively.

### Statistical Analysis

Data are presented as mean ± SD or mean ± SEM. Data were analyzed using GraphPad Prism software (Version 5, San Diego, CA). Groups were compared in a Bonferroni’s multiple comparison test (with selected pairs option) if ANOVA showed significant differences among groups. Student’s *t* test was applied to compare two experimental groups. *P* < 0.05 was selected as the criterion for statistical significance in all cases.

## RESULTS

### Ion Transport Mediated by DMT1: A Comparison of Transport Properties Inferred by Calcein Quench and Voltage Clamp Assays

Most previous pharmacological studies of DMT1 used a fluorescence quench assay with calcein as an indicator of intracellular iron. We have implemented this assay and, further, used voltage clamp techniques to verify that small molecules directly interacted with DMT1.

Consistent with previous studies using whole-cell voltage clamp techniques (Xu et al., 2004), changing the bath pH from 7.4 to 5.0 evoked an inwardly rectifying current in cells expressing hDMT1 (Fig. 1A). The inward current was increased by adding 2 μM Fe^++^ to the bath, but most of the current was still carried by protons. Substantially larger proton currents were measured when the bath pH was changed from 7.4 to 4.2 and the proton currents were hardly detectable at pH 6.0 (Fig. 1B). Expression of hDMT1 was controlled by an inducible promoter and no inward current was elicited by low pH in uninduced cells (Fig. 1C). This indicates that proton-conducting channels or transporters other than DMT1 do not significantly contaminate the voltage clamp measurements. Iron influx, as measured by calcein quench, is maximal at pH 6 (Fig. 1D), confirming previous reports using both calcein quench and ^55^Fe (Picard et al., 2000; Zhang et al., 2008).

**Fig. 1.**
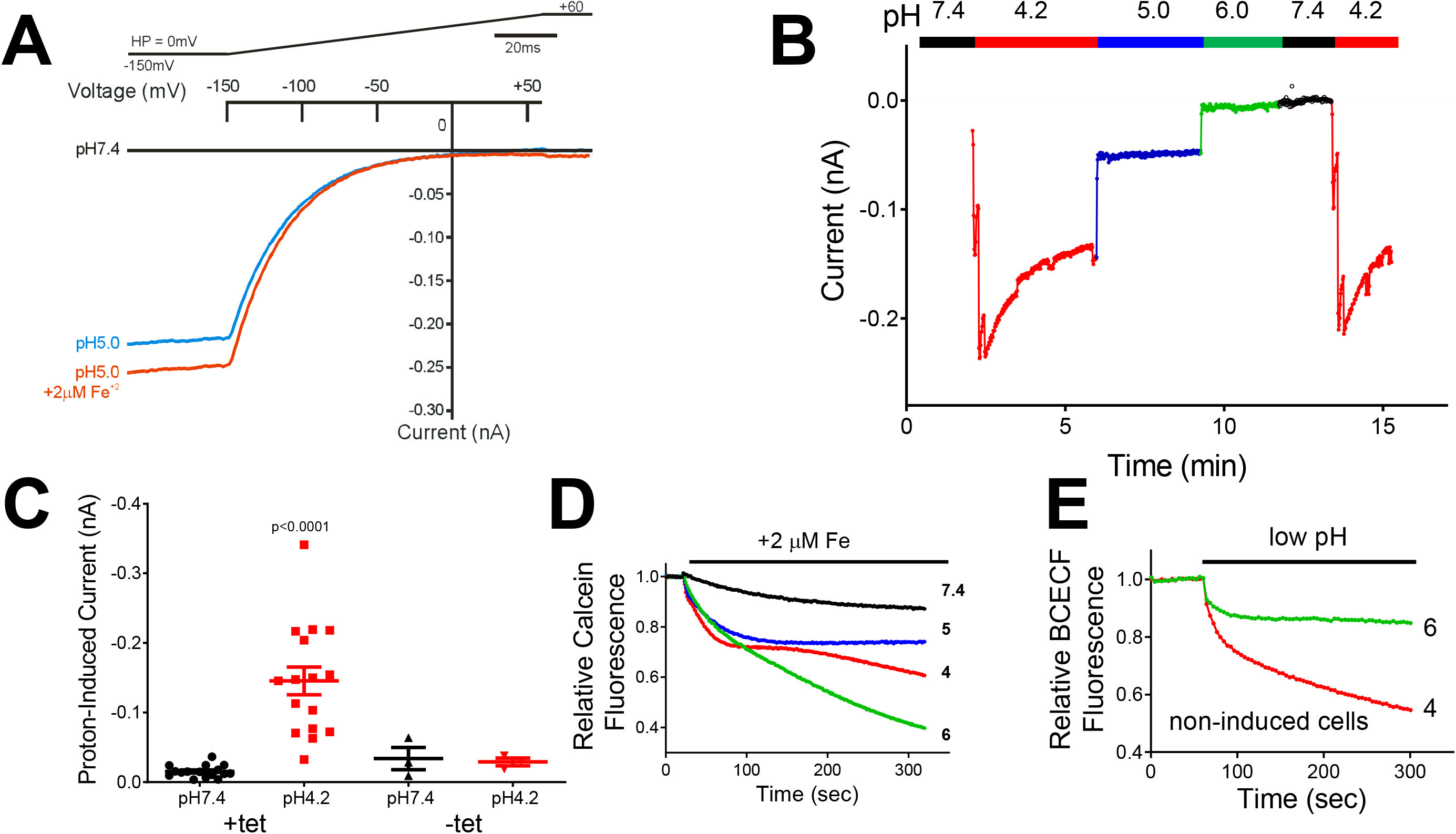
DMT1 transport reported by calcein quench and voltage clamp assays. (A) Current-voltage relationship for hDMT1. Whole-cell currents were elicited from a holding potential (HP) of 0 mV by repeated voltage ramps from −150 to +60 mV in 100 ms. Inward current was elicited by changing the bath pH from 7.4 to 5.0. A small additional current was elicited by adding 2 μM Fe^++^. (B) pH dependence of hDMT1 measured by voltage clamp. The solution exchanges are indicated above the current readings. Robust inward currents were elicited at −150 mV for pH 4.2 and pH 5 but not at pH 6. (C) Significant inward current at −150 mV was only observed in HEK cells at acidic pH when expression of hDMT1 was induced with Tet. This indicates the absence of significant endogenous proton conductance. (D) pH dependence of Fe^++^ transport measured by calcein quench assay. In contrast to panel B, influx is maximal at pH 6. (E) Rapid intracellular acidification in CHO cells when extracellular pH was lowered to 4. Intracellular pH was measured with BCECF, a pH indicator dye. The emission at 535 nm is plotted versus time.

In our voltage clamp studies the intracellular pH was unaffected by exposure to low pH bath solutions because the cells were dialyzed with a high concentration of pH buffer. In contrast, in the calcein quench assay exposure of cells to pH 4 caused rapid intracellular acidosis, as measured with the fluorometric dye BCECF (Fig. 1E). The intracellular acidosis quenches the fluorescence of calcein, reduces the quench due to iron binding (Tenopoulou et al., 2007) and also reduces the proton gradient that drives ferrous ion influx via DMT1. Exposure to low pH bath solution could also cause cell depolarization, further reducing the electrochemical driving force for iron influx. We thus conclude that the pH dependence of DMT1 transport is more reliably reported by voltage clamp studies and that DMT1-mediated current monotonically increases with the concentration of extracellular protons, although it is unclear whether influx of both protons and ferrous iron increase in parallel.

#### Synthesis of Potent Multimeric DMT1 Inhibitors

We used a calcein quench assay to screen more than 175,000 compounds and identified several lead series, some of which have been reported previously (Cadieux et al., 2012; Zhang et al., 2012). We synthesized analogs of the original leads and optimized for compounds with good potency, low systemic bioavailability and in vivo efficacy, as outlined in Fig. 2. Both the calcein quench and voltage clamp measurements were used to confirm block of DMT1 transport in vitro, and a Caco-2 epithelial transport assay evaluated expected in vivo systemic uptake (Artursson et al., 2001). Inhibition of iron absorption was evaluated in an acute model of iron hyperabsorption in anemic rats, as described below.

**Fig. 2.**
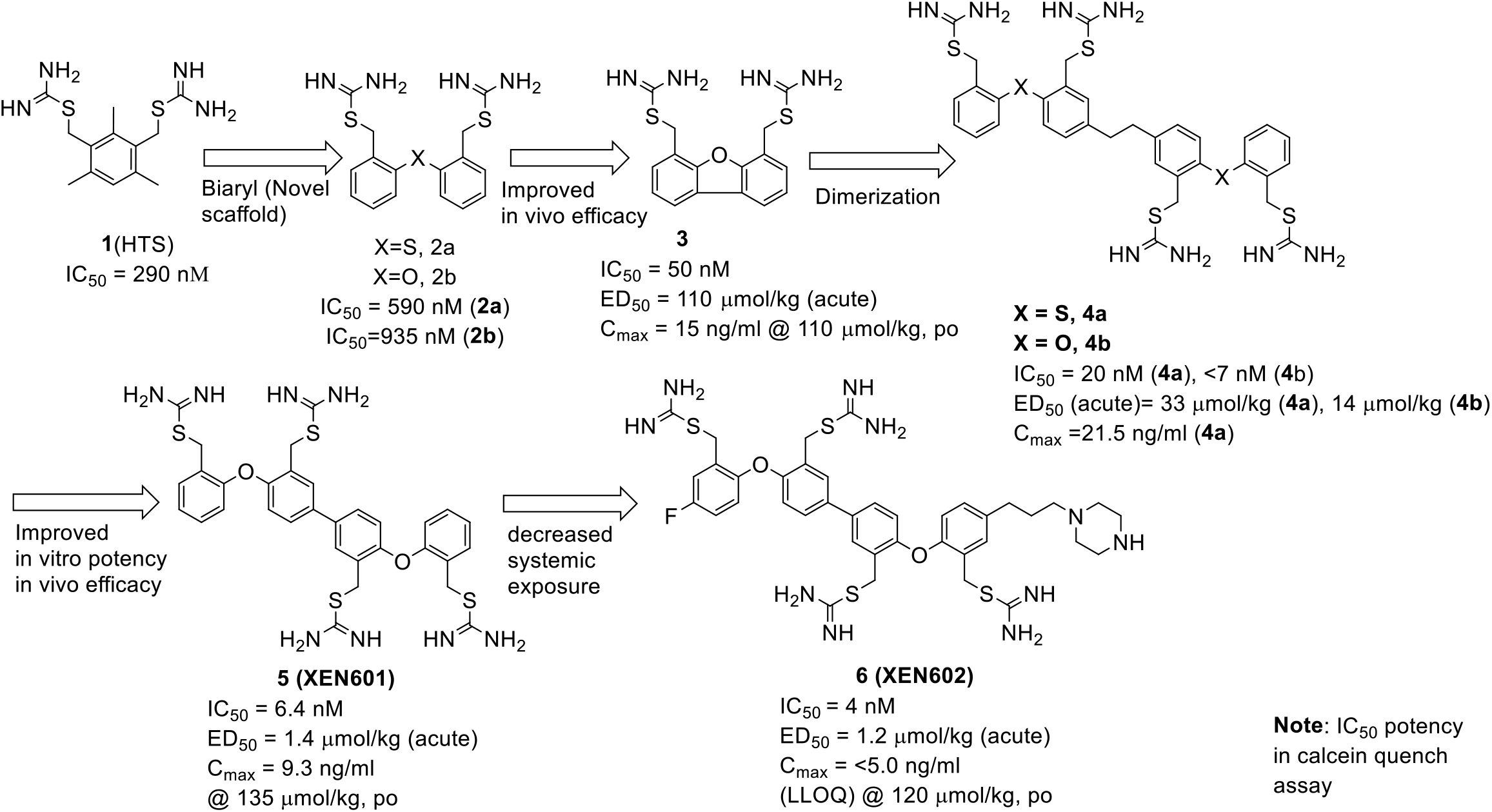
Lead optimization of DMT1 inhibitors to generate XEN601 and XEN602. Compound **1** was identified in a high-throughput screen (HTS). The progression to optimized monomers and generation of dimers is indicated schematically. LLOQ, lower limit of quantitation.

The lead compound **1** had modest potency and Caco-2 permeability (Papp = 0.9 × 10^6^ cm/s [a→b], 1.2 × 10^6^ cm/s [b→a]) (Zhang et al., 2012). As reported previously, a medicinal chemistry effort led to novel structures, a biaryl ether and tricyclic dibenzofuran *bis*isothiourea, compounds **2** and **3** (Fig. 2). Compound **2b** demonstrated significant in vivo activity in the acute iron hyperabsorption model. Compound **3** had greater potency, both in vitro and in vivo. Compound **3** was well tolerated with no clinical signs at a dose of 871 μmol/kg but had significant oral exposure at the ED_50_ of 110 μmol/kg (C_max_ = 15 ng/ml).

Although these compounds demonstrated an encouraging correlation between IC_50_ for inhibition of DMT1 transport and ED_50_ in an animal model of iron overload, with no acute toxicity, the efficacious doses and systemic exposures were higher than acceptable for therapeutic development. We successfully improved potency and lowered systemic exposure by generating multimers. The dimer, compound **4a**, with a *tetra*isothiourea moiety, was generated by linking the compound **2a** monomers. Although we expected that improved efficacy would only be obtained with longer linkers, compound **4a** showed 30-fold greater potency in both the calcein quench and voltage clamp assays (IC_50_ = 20 nM and 29 nM, respectively) as compared to the corresponding monomer (IC_50_ = 590 and 2140 nM, respectively). Compound **4a** showed significant inhibition of iron absorption when evaluated in an acute model of iron hyperabsorption in anemic conscious rats (ED_50_ = 33 μmol/kg, as described below) and was well tolerated with no clinical observations up to a maximum tolerated dose of 1065 μmol/kg. However, the systemic exposure of compound **4a** at the ED_50_ dose (C_max_ = 21.5 ng/ml, p.o.) was still unacceptably high. The oxygen analog, **4b**, showed in vitro potency and in vivo efficacy similar to that of **4a**.

In order to achieve in vivo efficacy at even lower doses and lower systemic exposures, we explored the chain length of the dimer. We found that the optimal dimer is one with a direct bond between the aryl moieties, compound **5 (XEN601)**. This molecule showed excellent in vitro potency, with an IC_50_ = 6.4 nM. In the acute model of iron hyperabsorption, **XEN601** had an ED_50_ =1.4 μmol/kg. **XEN601** was well tolerated with no clinical observations up to a maximum tolerated oral dose of 579 μmol/kg but showed a significant level of systemic exposure after oral administration (C_max_ = 9.3 ng/ml at 135 μmol/kg, p.o.).

Also, **XEN601** is a potent inhibitor of CYP3A4 (IC_50_ < 1 μM, data not shown), which is undesirable because CYP3A4 is abundant in the gut (Rommel and Richard, 2017).

In addition to the dimeric constructs, we tried to achieve high potency and low systemic exposure by introducing heteroatoms into the biaryl ether moiety or by adding polar side chains to the monomers (see Supplemental Table S1).

The greatest success in retaining potency while reducing systemic exposure was obtained by addition of a large polar side chain to **XEN601**. Adding an alkyl tail with a tertiary or quaternary amine was tolerated but was not an improvement over **XEN601** (compounds **17** and **18**; see Supplemental Table S1). However, adding a larger piperazine ring with an alkyl chain linker proved very successful (compound **6**, also known as **XEN602**). This compound, as well as variants with a polyethylene glycol side chain (compounds **20** and **21**; see Supplemental Table S1) retained the potency of **XEN601** but had much lower systemic exposure. For example, **XEN602** has bioavailability in rats of 0.1% (C_max_ = 320 ng/ml) at an oral dose of 240 μmol/kg, which is more than 150 times higher than the ED_50_. Moreover, the compound was rapidly cleared from systemic circulation, with a half-life of 0.55 h. Based upon these results, **XEN601** and **XEN602** were characterized as potential clinical candidates, as described below.

#### Block of Proton and Divalent Cation Transport via DMT1

For our pharmacological studies, we utilized both (a) calcein quench measurements at pH 6 to measure influx of Fe^++^ and (b) voltage clamp studies at bath pH 5.0 or 4.2, in which most of the current was carried by protons. In some experiments, 30 μM Mn^++^ was added to quantify current carried by divalent cations. In general, the calcein quench assay indicated a lower IC_50_ than that found by voltage clamp studies. An example of the voltage clamp studies is shown in Fig. 3A. These studies revealed much greater ionic transport at pH levels encountered in the gut than previously inferred from fluorescence assays alone. Block by **XEN601** is quantified by measuring the reduction in inward current elicited by a bath solution at pH 5.0. As also seen in Fig. 1, the absence of outward current at +60 mV indicates that linear leak current is negligible and that essentially all of the current is carried by protons via DMT1.

**Fig. 3.**
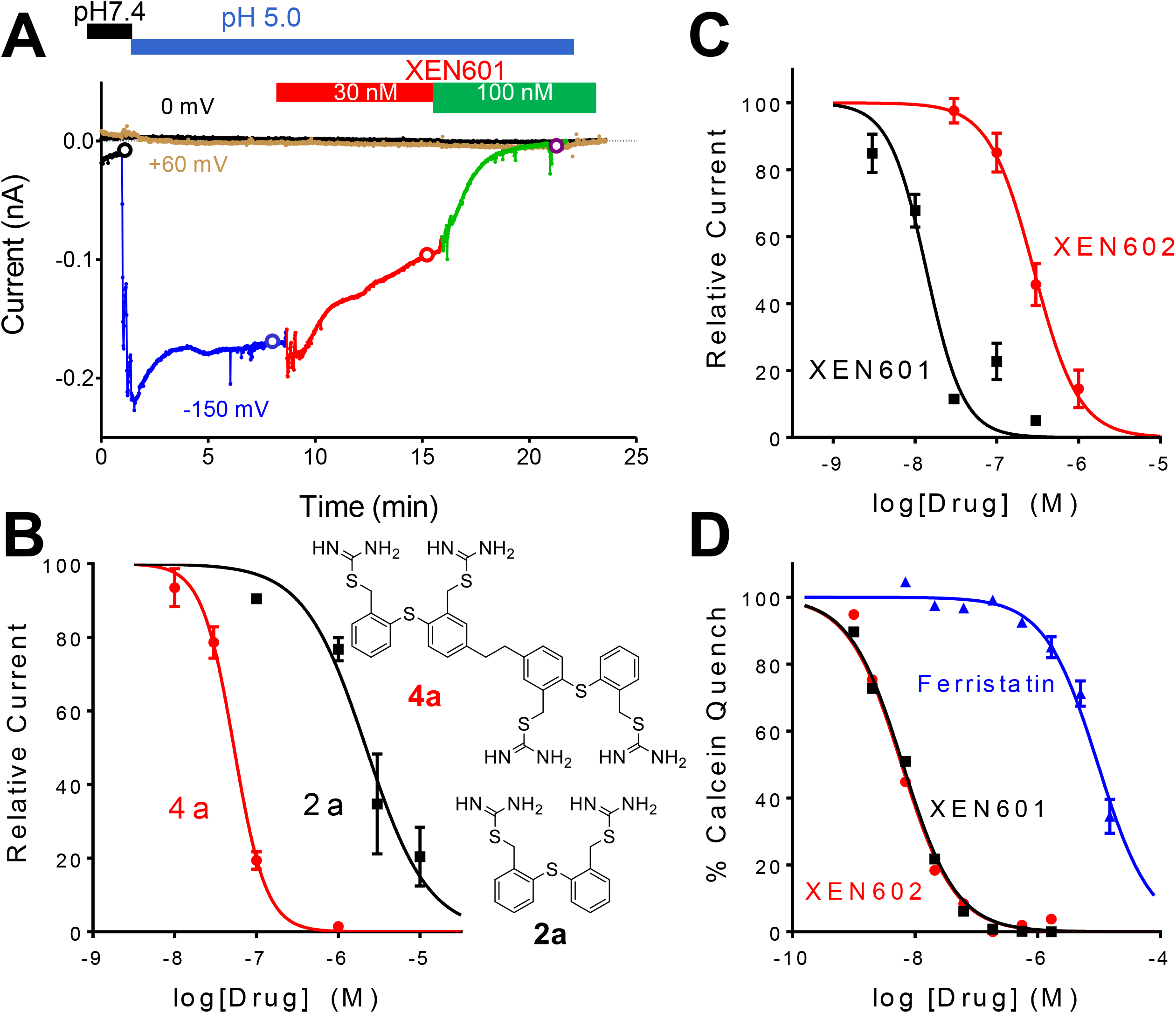
In vitro characterization of DMT1 inhibitors XEN601 and XEN602. (A) Voltage clamp analysis of effects of **XEN601** on DMT1-mediated proton transport. The solution exchange protocol is indicated above the current readings. Upon transitioning from pH 7.4 to pH 5.0, an initial spike of inward current quickly decayed until it reached steady-state after approximately 5 min. Thereafter, subsequent addition of 30 nM and 100 nM **XEN601** at pH 5.0 showed concentration-dependent inhibition of proton current. (B–D) Concentration-response curves for inhibition of DMT1. (B) Comparison of DMT1 block by monomeric and dimeric constructs. The dimer (**4a**) was approximately 40-fold more potent than the monomer (**2a**) in blocking DMT1-mediated proton transport at pH 5.0 measured by voltage clamp (IC_50_ = 53 nM and 2.2 μM, respectively). (C) Concentration dependence of block of hDMT1 by **XEN601** and **XEN602** measured by voltage clamp (IC_50_ = 14 nM and 280 nM, respectively). Proton currents were measured at pH 5.0 and at −150 mV. (D) Concentration dependence of block measured by the calcein quench assay at pH 6.0. **XEN601** and **XEN602** were approximately equipotent (IC_50_ of 6.4 nM and 4.0 nM, respectively) and much more potent than ferristatin (IC_50_ = 9.6 μM). Block by both compounds and by ferristatin was well fit with a Hill coefficient of 1.0.

Compound **4a** was studied in detail by voltage clamp methodology because it was the first compound to demonstrate the benefit of a dimeric construct (Fig. 3B). As for most of our studies, currents were measured at pH 5.0 without the addition of Fe^++^ or Mn^++^. The monomer equivalent, compound **2a**, produced block that was best fit with a Hill coefficient of 1.1 and IC_50_ = 2.2 μM. In contrast, compound **4a** was about 40-fold more potent (IC_50_ = 53 nM) and the concentration dependence of block was fit with a Hill coefficient of 2.1. This suggests that compound **2a** binds with a 1:1 stoichiometry to produce block whereas compound **4a** binds with a 2:1 stoichiometry.

The concentration dependence of inhibition of DMT1-mediated currents by **XEN601** and **XEN602** measured by voltage clamp (Fig. 3C) was best fit by Hill coefficients of about 2, as seen for compound **4a** (Fig. 3B). For **XEN601** and **XEN602**, the Hill coefficients were 1.9 and 1.6 when protons were the charge carrier at pH 5. Both compounds were very potent inhibitors of ferrous ion influx in the calcein quench assay (Fig. 3D). Since the sensitivity of this assay depends upon the level of expression of DMT1, we also quantified block by a well-documented pharmacological agent, ferristatin (NCI306711). This compound was the most useful DMT1 inhibitor identified in our screening campaign using a calcein quench assay; the IC_50_ in our studies is similar to that reported previously (Buckett and Wessling-Resnick, 2009). In our voltage clamp studies, ferristatin was inactive at 30 μM (data not shown). The antioxidant ebselen also reportedly blocks iron uptake via DMT1 (Wetli et al., 2006), but we found that it was inactive in the calcein quench assay at 15 μM and by voltage clamp at 1 μM (data not shown). Thus, **XEN601** and **XEN602** are >1000-fold more potent than previously available inhibitors.

The specificity of **XEN601** and **XEN602** was evaluated in a panel of radioligand binding assays (see Supplementary Table S2). Both compounds inhibit binding to multiple receptors with cationic ligands. Thus, both compounds are potentially pleiotropic and compound safety is conferred by (and dependent upon) low systemic availability.

### In Vivo Characterization of DMT1 Inhibitors

#### Development of an Acute Iron Challenge Model in the Rat and Characterization of XEN602

We wished to develop a simple in vivo assay that would indicate the efficacy of test compounds for reducing iron uptake from the duodenum via inhibition of DMT1. DMT1 is initially expressed at low or undetectable levels in samples of normal rat duodenum analyzed by western blotting (data not shown). We placed rats on an iron-deficient diet (2 ppm iron) for 4 weeks to induce a state of iron deficiency. At this point, measurements of Hb levels in blood indicated that the rats were severely anemic (Hb levels reduced from 14 to 6 g/dl; data not shown), and samples of duodenum analyzed by western blot showed a prominent band of DMT1 protein at 55 kDa (data not shown). An oral dose of 1 mg/kg of iron gave a 20- to 30-fold elevation in plasma or serum iron 1 h after dosing, and we subsequently used this dose as our standard acute iron challenge. We observed similar results for **XEN601** and **XEN602** in this model and, thus, only data for **XEN602** are reported here.

We used this assay to examine whether dimeric constructs have enhanced activity over the monomer subunits, similar to the enhancement of inhibition of DMT1 observed in vitro. **XEN602** is formed from two dissimilar monomers and we determined the effect of each monomer and of **XEN602** on serum iron levels after challenge (Fig. 4A). The compounds were administered 1 h prior to iron challenge. Under these conditions, a dose of 23 μmol/kg **XEN602** blocked approximately 95% of the increase in serum iron seen in untreated controls (*P* < 0.001 compared to vehicle). Even at a 2-fold higher dose, one of the component monomers of **XEN602**, the 3-fluoro analog of compound **2b**, was approximately 25-fold less potent than **XEN602**. At an almost equivalent dose the other component monomer of **XEN602**, compound **9**, was approximately 8-fold less potent than **XEN602** (*P* < 0.01).

**Fig. 4.**
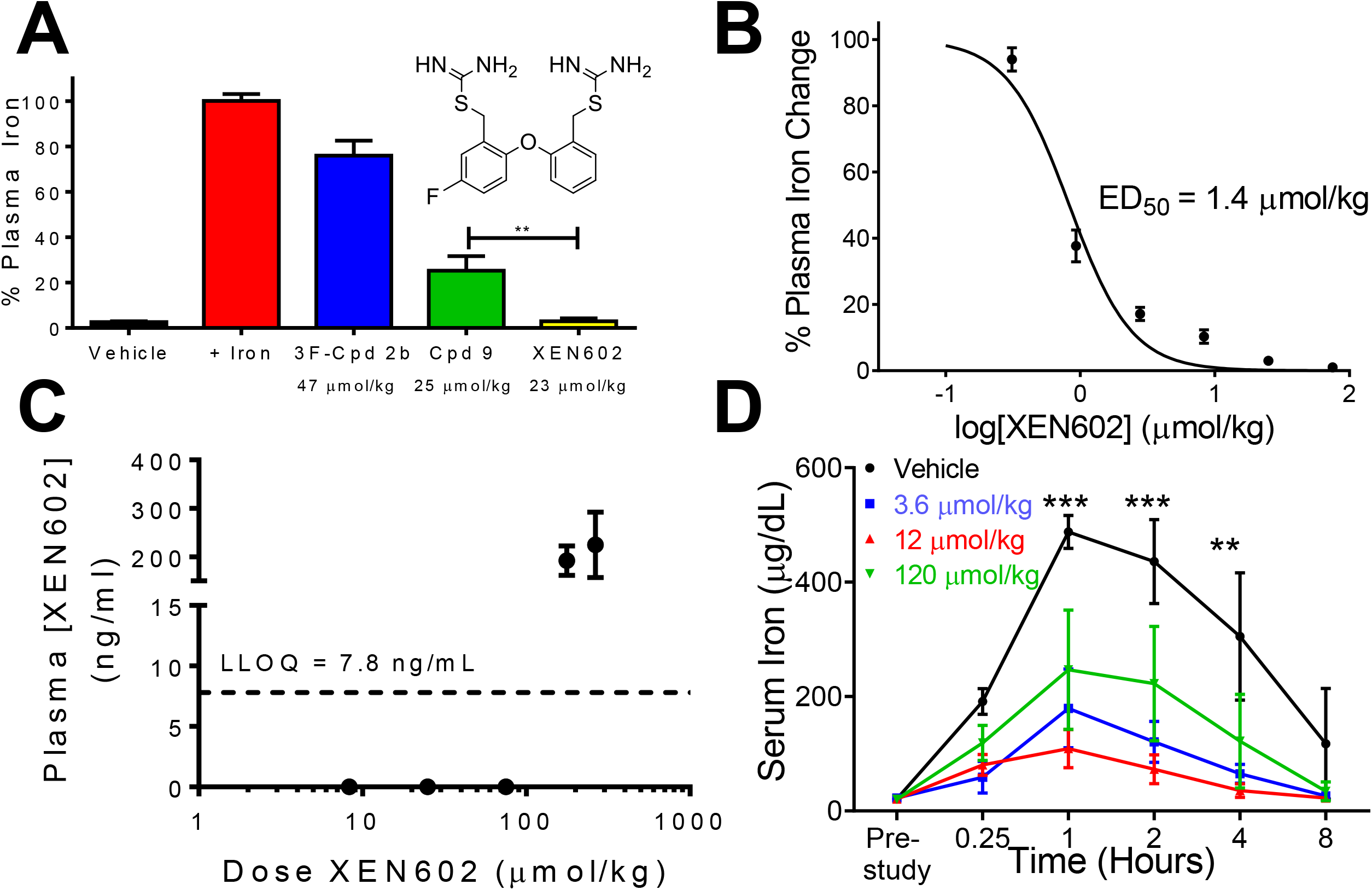
Effects of single doses of XEN602 in an acute iron challenge model in rats and weanling pigs. (A) **XEN602** exhibits superior potency to its component monomers 3-fluoro-compound **2b** (structure shown in the inset) and compound **9** in an acute iron challenge model in the rat. Doses were based on molarity to account for substantial differences in the molecular weight of the compounds. (B) The increase in serum iron determined in rats 1 h after a 1 mg/kg oral iron challenge is blocked in a dose-dependent fashion by a single oral dose of **XEN602**. (C) Observed plasma concentrations of **XEN602** at 1 h post dose. (D) The increase in serum iron determined in weanling pigs from 0.25 to 8 h after a 3 mg/kg oral iron challenge is blocked in a dose-dependent fashion by a single oral dose of 3.6 and 12 μmol/kg **XEN602**. The effects of the compound were observed up to 4 h after dosing. \*\**P* < 0.01, \*\*\**P* < 0.001, comparison between vehicle and 12 μmol/kg dose groups by two-way ANOVA.

We generated a dose-response curve for **XEN602** in this model (Fig. 4B). Doses between 0.31 and 75 μmol/kg were tested, enabling calculation of an ED_50_ of 1.4 μmol/kg (1.2 mg/kg).

Measurements of plasma levels of **XEN602** performed at termination (1 h following dosing) at dose levels up to 75 μmol/kg demonstrated that, in all but one experiment, compound was undetectable, with a lower limit of quantitation of 7.8 ng/ml (Fig. 4C). Thus, **XEN602** demonstrated a potent block of iron uptake mediated by DMT1 in vivo with negligible compound levels observed in plasma, suggesting the activity of the compound was successfully restricted to the gut at a dose ~100-fold higher than the ED_50_. However, in one of four animals dosed at 23 μmol/kg, a plasma level of 117 ng/ml was observed (data not shown). We postulated that the high systemic exposure was due to a compromised GI permeability barrier. This idea was tested by co-administering ibruprofen at ≤200 mg/kg, and we observed similar variability in plasma concentration (data not shown).

#### Development of an Acute Iron Challenge Model in the Weanling Pig and Characterization of XEN602

Laboratory rodents have a lower proportion of heme iron in their diet than most large mammals, leading to concern that inhibition of DMT1 might be of greater consequence in rats. Thus, we established a similar anemia model in weanling pigs and tested the efficacy of **XEN602** in a similar fashion as described for the rat. Neonatal pigs are normally given a dose of 200 mg of iron dextran shortly after birth to prevent anemia. Therefore, in order to develop a model of mild anemia, neonatal pigs were given a reduced dose of 50 mg of iron dextran at birth. By 4 weeks of age these animals developed a mild to moderate anemia (average Hb levels were 7.83 ± 1.77 mg/dl, compared with a normal range of 10–16 mg/dl).

A 3 mg/kg iron challenge was an optimal dose in this model and produced a >20-fold increase in serum iron levels 1 h following an iron challenge in vehicle-treated animals (21 ± 5 vs. 488 ± 50 μg/dl, n = 3). Doses of 3.6, 12, and 120 μmol/kg **XEN602** were then given p.o. 1 h prior to the iron challenge, and plasma samples were collected for serum iron measurement 0.25 to 8 h following iron challenge (Fig. 4D). A dose-dependent inhibition of iron uptake was observed at 3.6 and 12 μmol/kg, but less inhibition was observed at 120 μmol/kg. The reasons for the less efficacious response observed in the 120 μmol/kg versus the lower dose groups are unclear. In addition, the dose-response curve did not appear to be as steep as that observed in the rat. Data from this and subsequent experiments (data not shown) suggested that the ED_50_ for **XEN602** in this model is between 3.6 and 12 μmol/kg. Thus, we were able to demonstrate that **XEN602** was efficacious in an acute setting in a large mammal at pharmacologically viable doses.

In this experiment, we also determined plasma concentrations of **XEN602** 0.25 to 8 h following iron challenge. Using a bioanalytical method with a lower limit of quantitation of 10 ng/ml, no compound could be detected in plasma at these time points for doses up to 3.6 and 12 μmol/kg. Two of three animals in the 120 μmol/kg dose group absorbed detectable **XEN602** into their bloodstream, with C_max_ values of 1221 and 677 ng/ml 15 min following iron challenge. Thus, at doses up to 3-fold greater than the estimated ED_50_, **XEN602** was undetectable in plasma (data not shown).

#### Development of a Subchronic Iron Challenge Model and Characterization of XEN602

We also sought to establish a more physiological iron challenge regime, both to try to model the physiological conditions under which therapeutic DMT1 inhibitors might be effective, and for use in chronic efficacy testing of compounds that antagonize DMT1 function. We induced anemia, iron deficiency and DMT1 upregulation as before, by placing rats for four weeks on an iron-deficient diet. Anemic rats were maintained on a low-iron diet and gavaged daily with 0.3 mg/kg ferric citrate 1 h following a single daily dose of vehicle or oral **XEN602** at either 3 or 10 μmol/kg. Recovery from a state of iron deficiency was monitored by testing Hb on a weekly basis (Fig. 5A). Under this regime, vehicle-treated animals progressively recovered increased Hb levels over 17 days, but the Hb levels of rats dosed with **XEN602** remained unchanged, indicating effective and maintained block of iron uptake under these conditions. A dose-related effect on weight gain was observed (Fig. 5B), but as no effect on food consumption was observed (data not shown) and no impairment of weight gain was observed in non-anemic animals at this dose (Fig. 6A), the effect on weight gain is most likely a secondary consequence of maintaining an anemic state in these animals. Similar results were obtained in a variation of this model where the recovery phase included access to an iron-replete diet for 3 h per day following compound dosing instead of ferric citrate by gavage (data not shown).

**Fig. 5.**
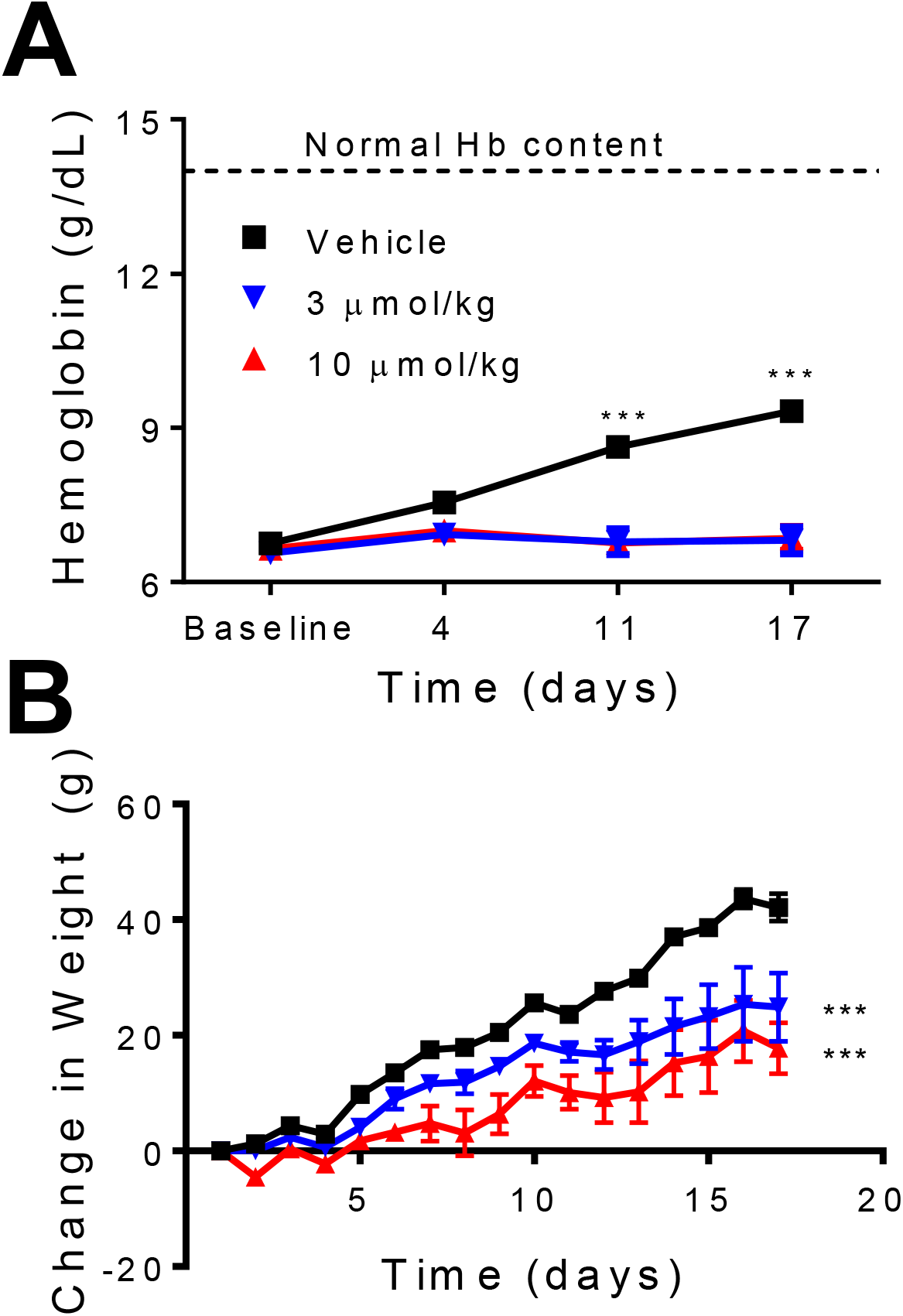
Effects of repeat doses of XEN602 in a subchronic iron challenge model in the rat. (A) Hemoglobin levels in anemic rats were maintained on a low-iron diet with daily gavage of iron (0.3 mg/kg ferric citrate) 1 h following a single daily dose of vehicle or oral **XEN602** at either 3 or 10 μmol/kg. (B) Reduced weight gain in rats dosed daily for 17 days with 3 or 10 μmol/kg **XEN602**. *** *P* < 0.001, comparison between vehicle and 10 μmol/kg dose groups by two-way ANOVA.

**Fig. 6.**
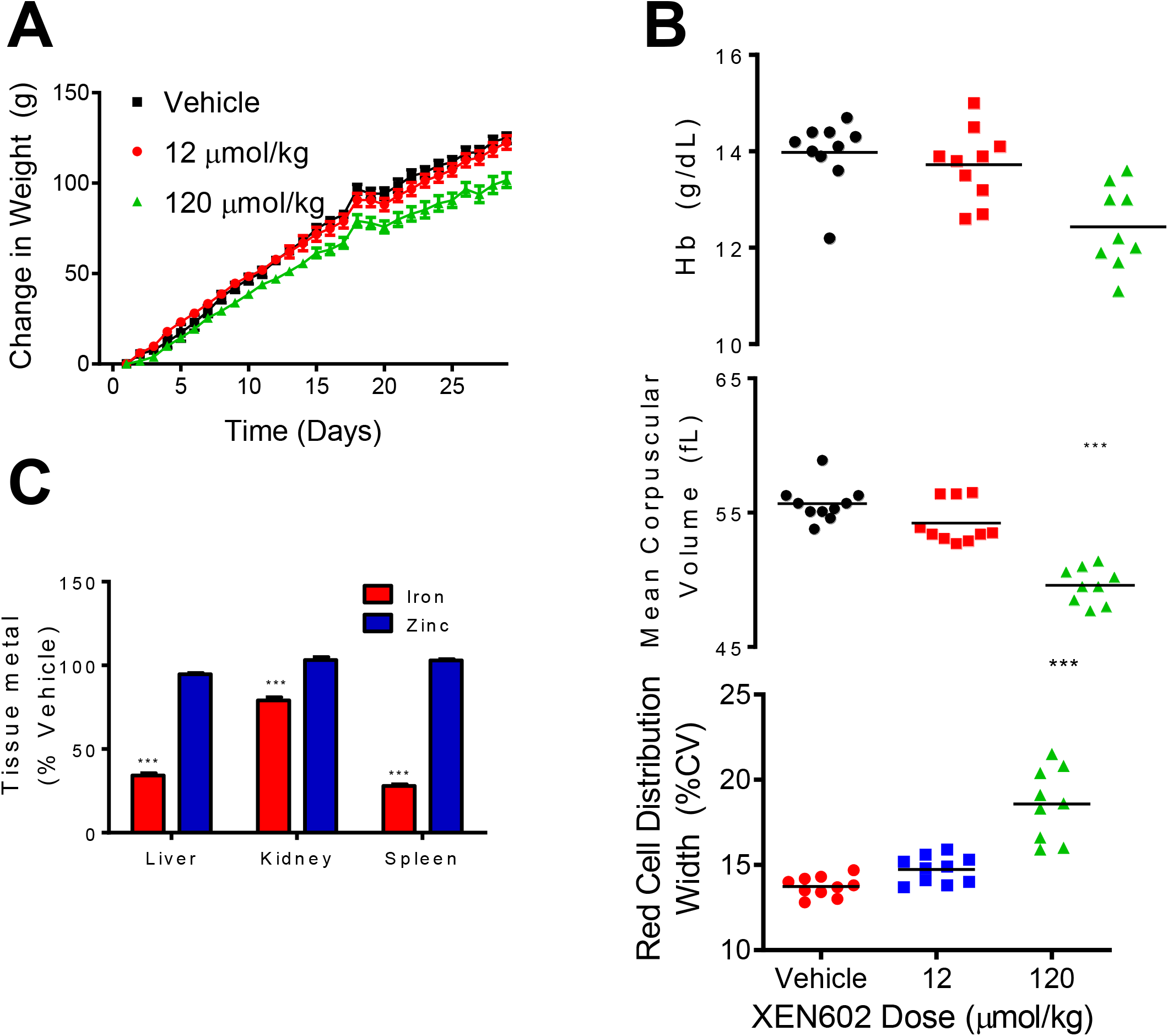
Safety assessment studies in non-anemic Sprague Dawley rats: Increased permeability in compromised GI tract and consequences for drug safety. (A) Reduced weight gain in rats dosed daily for 28 days with 100 mg/kg **XEN602**. (B) Reduced hemoglobin, reduced mean corpuscular volume and increased red cell distribution width in terminal blood samples from rats dosed daily for 28 days with 100 mg/kg **XEN602**. (C) Reduced iron, but not zinc, in liver, kidney and spleen in terminal tissue samples from rats dosed daily for 28 days with 100 mg/kg **XEN602**. All data presented as mean + SEM. \*\*\**P* < 0.001 when compared to vehicle group.

From these experiments we conclude that **XEN602** is efficacious in vivo in blocking the uptake of iron in the gut after an acute iron challenge in both the rat and weanling pig. The compound also blocks effective iron uptake, as assessed by maintenance of anemia, in iron-deficient rats allowed limited access to iron. This indicates that the mechanism does not desensitize over a time frame of at least 2 weeks.

#### Safety Assessment

##### Repeat dose study

**XEN602** was evaluated using a once-a-day repeat oral dosing regimen of 0, 12, and 120 μmol/kg for 28 days in healthy (non-anemic) Sprague Dawley rats maintained on an iron-replete diet. The primary findings were confined to the high-dose group where body weight gain was significantly lower (18% and 17%) compared to the vehicle and low-dose groups, respectively (Fig. 6A). The lower body weight correlated with ~10% lower cumulative food intake during the course of the study in the high-dose group (data not shown). Gut motility was unaffected by **XEN601** at a dose of 120 μmol/kg (Supplementary Fig. S1), and it is unlikely that **XEN602** has an effect on transit time. Hematologic analysis of terminal blood samples showed decreases in mean corpuscular volume and Hb concentration as well as an increase in red cell distribution width in the high-dose group (Fig. 6B). These findings are consistent with the presence of an iron-deficiency anemia related to compound block of DMT1-mediated iron absorption. Further support for a state of iron deficiency in the 120 μmol/kg group was a complete lack of iron staining in spleens from these animals (data not shown). Moreover, the iron concentration in liver, kidney and spleen in the high-dose group was significantly decreased (Fig. 6C; low-dose group not analyzed).

Since divalent metals other than iron can be transported by DMT1, we evaluated the tissue content of multiple metals under conditions in which iron transport was greatly reduced in our rodent efficacy model. These studies were particularly important because **XEN602** is a polycation that might inhibit other cation transporters in the gut. However, polycations zinc (Fig. 6C), copper, calcium, cadmium, magnesium, manganese, nickel and barium were generally unaffected in these tissues, but cobalt was elevated in the spleen of rats dosed at 120 μmol/kg (data not shown). Thus, **XEN602** is a highly selective inhibitor of DMT1 in the gut and inhibition of DMT1 can be achieved without compromising the uptake of other divalent cations, at least over the time course of this study.

Variable systemic exposure was observed in this study; that is, for animals dosed at 120 μmol/kg, mean C_max_ was 26 ± 32 ng/ml, ranging from 7 to 110 ng/ml. Based upon the body weight loss and elevated transaminases at 120 μmol/kg /day (data not shown), the no-adverse-effect level (NOAEL) was considered to be 12 μmol/kg /day, but it is likely between 12 and 120 μmol/kg /day.

##### Maximum tolerated dose

As described above, in efficacy studies **XEN602** usually had low oral bioavailability. However, higher exposure was sometimes observed in a few animals, and there was a risk that higher exposures could be encountered in cases of compromised GI permeability barriers, such as in humans with gastric or duodenal inflammation or ulceration. To better understand the consequences of systemic exposure, we increased exposure either by using very high oral doses of drug or by IV infusion. **XEN602** was administered orally to rats in doses of 240, 360, 480 μmol/kg. At doses >240 μmol/kg, serious adverse effects were noted, including skin erythema followed by cyanosis, salivation, labored breathing, marked respiratory depression, hemoconcentration and elevated clotting times (aPTT and PT). Plasma levels were variable, particularly in the 360 μmol/kg group, where C_max_ values ranged from 172 to 4598 ng/ml, perhaps reflecting saturation of an efflux transporter in the GI tract. Similar signs were observed with a single IV bolus injection (1.2 or 0.6 μmol/kg), a constant IV infusion (0.5 μmol/ml; 0.74 ml/h) or a single intraperitoneal administration (6 or 60 μmol/kg). The IV infusion study facilitated more accurate characterization of the concentration-effect relationship. Plasma levels up to 200 ng/ml were generally well tolerated, but early clinical signs were observed at higher concentrations, typically including skin erythema, irregular or increased breathing and bluish extremities. Plasma levels of approximately 400 ng/ml were associated primarily with cyanosis. At concentrations ≥1000 ng/ml, more severe signs became apparent, with euthanasia frequently being indicated.

To evaluate whether these effects were species specific, a subsequent IV study was performed in domestic pigs. With the exception of clotting time changes, the pigs showed signs similar to rats with the following additional abnormalities documented: vomiting, diarrhea, hypotension, cardiac electrical disturbances and convulsions. These effects were noted after single IV doses of 0.12 or 0.36 μmol/kg. Intravenous doses of 0.012 and 0.036 mg/kg, on the other hand, were generally well tolerated, indicating a steep dose-response relationship in pigs as in rats. Plasma levels associated with adverse effects were in the same range as in the rat IV study. There was no evidence of methemoglobin formation, and the mechanism of toxicity is unknown.

## DISCUSSION

We report novel inhibitors of DMT1 that are substantially more potent than previously reported compounds (Brown et al., 2004; Buckett and Wessling-Resnick, 2009). We achieved this potency by linking monomers of much less potent compounds and thereby also greatly enhanced localization to the site of action in the duodenum. The lead compound, **XEN602**, is a remarkably selective inhibitor of dietary iron uptake with little effect on the transport of other cations and is therefore a powerful pharmacological tool for characterizing the role of DMT1 in dietary iron absorption.

### Calcein Quench

The calcein quench assay is an exquisitely sensitive measure of iron uptake and can be readily adapted to high-throughput screening methodologies (Breuer et al., 1995; Picard et al., 2000). However, multiple mechanisms of action other than inhibition of DMT1 can result in inhibition of calcein quench. In particular, co-transport of Fe^++^ and protons is electrogenic and is diminished by depolarization of the membrane (Gunshin et al., 1997), so that some compounds that diminish calcein quench could do so by depolarizing the cells that heterologously express hDMT1 rather than acting directly on the transporter. In addition, compounds that alter the redox status of the cell can register as inhibiting calcein quenching because the quench is due to binding of ferric ions (Thomas et al., 1999), whereas DMT1 transports ferrous ions.

### Biophysics of DMT1

To our knowledge, this is the first report that utilizes both the calcein quench and voltage clamp assays to quantify transport via DMT1. These assays provide complementary information on transport in that the fluorescence assay measures iron transport with great sensitivity, while the voltage clamp technique measures total ionic transport under conditions where the ionic gradients can be maintained. Intracellular dialysis to achieve heavy buffering of pH reveals that proton transport does not readily saturate at an extracellular pH of 6 and that protons carry most of the current at lower pH. Protons can be transported without Fe^++^. Block of proton currents indicates that **XEN601** and **XEN602** directly inhibit the transporter, rather than chelating divalent cations. We found no evidence that Fe^++^ can be transported without an inward pH gradient, in contrast to a study with DMT1 expressed in oocytes (Mackenzie et al., 2006). However, small current components can be detected more readily in oocyte experiments and it is possible that a small current went undetected.

Voltage clamp measurements allow for complete control of the ionic composition on both sides of the membrane and thus are better suited to establish a direct effect on the transporter. In addition, the assay provides a linear measure of transport rate. However, very high levels of protein expression are required in order to establish a robust assay. Moreover, many cell lines commonly used for heterologous expression also express an endogenous proton conductance that potentially confounds measurements of transport via DMT1.

Previous voltage clamp studies suggested a pH dependence of transport that is dramatically different from that inferred from calcein quench or ^55^Fe influx assays (Picard et al., 2000; Zhang et al., 2008): the influx assays indicate that influx is optimal at about pH 5.5–6, whereas voltage clamp studies indicate that proton currents increase with the external proton concentration up to at least pH 4 (Xu et al., 2004). Likewise, in oocytes the influx of iron mediated by DMT1 under voltage clamp increases monotonically up to at least pH 5.2 (Mackenzie et al., 2006). It is advantageous to quantify block of DMT1 transport at very low pH in order to make the currents as large as possible, but we needed to establish that these currents were entirely due to DMT1 and therefore needed to account for the apparent inconsistency with calcein quench studies.

In order to engage adjacent DMT1 transporters with multimers, we anticipated using linkers of about 10 nm (10–20 kDa), as found in a study of CFTR transporters in the GI tract (Sonawane et al., 2008). Instead, we found that very short linkers are most effective. The increase in potency seen with a dimer is not simply a function of greater charge density because compound **19** has less charge and similar potency to **XEN601** and **XEN602**. The X-ray crystal structure of a homologous bacterial transporter (ScaDMT) indicates a protein fold and two similar subdomains in each monomer (Ehrnstorfer et al., 2014), and a similar structure is inferred by bioinformatic analysis of the SLC11 family (Cellier, 2012). This symmetry suggests two similar binding sites in a monomer rather than a dimeric structure or binding that spans two nearby monomers.

### Physiological Role of DMT1 in Rats and Large Animals

We observed a good correlation between in vitro potency for block of DMT1 and potency in the acute iron absorption assay. With **XEN601** and **XEN602** we were able to completely inhibit iron uptake without effect on the uptake of other divalent cations. Thus, we were able to pharmacologically mimic the effect of DMT1 knockout specific to the duodenum. These results suggest that DMT1 is the major pathway for uptake of non-heme iron, but it is possible that **XEN601** and **XEN602** block additional unidentified transporters. However, intestinal iron uptake is also greatly diminished by loss-of-function mutations in DMT1 in Belgrade rats and microcytic anemia (*mk*) mice (Fleming et al., 1997; Fleming et al., 1998), which lends support to the idea that the inhibition of iron absorption by our compounds is due entirely to block of DMT1.

We present the first studies of DMT1 block in a non-rodent species. Previous genetic and inhibitor in vivo studies of enterocyte DMT1 have been conducted with rodents, but these animals have a diet that is very low in heme-containing protein. Thus, DMT1 function might differ in a larger animal with a more varied diet. **XEN602** appeared to be slightly less potent in the pig as compared to the rat (estimated ED_50_ of 1.2-3.6 μmol/kg vs. 1.4 μmol/kg). However, we had less opportunity to optimize the weanling pig model, and the modest difference in ED_50_ may reflect lesser refinement of assay conditions. In both cases, the iron challenge is greater than would be encountered clinically, and it is encouraging to see efficacy at pharmacologically reasonable doses in two species. Moreover, the compounds showed good efficacy in a subchronic iron uptake assay in the rat, indicating effective and maintained block of iron uptake for at least 17 days with no evidence of desensitization of inhibition.

### Development of Putative Gut-Restricted Therapeutics

In many respects, we achieved our therapeutic goal with **XEN602** in that we could completely inhibit iron uptake at a low dose of drug, yet have negligible systemic exposure at a 100-fold higher dose. However, this compound has serious off-target activities that require negligible systemic exposure and a steep dose dependency. Our concern regarding exposure when the GI tract is compromised prompted us to perform experiments using non-steroidal anti-inflammatory drugs to disrupt the intestinal barrier, but we were unable to achieve a consistent effect on plasma drug levels. Moreover, we observed unpredictable substantial systemic exposure both in rats (described above) and pigs. This unpredictability of systemic exposure coupled with the steep concentration-response relationship culminating in mortality led us to halt development of the compound.

#### Prospects for Therapeutically Useful DMT1 Inhibitors

We can suggest several strategies to utilize the current chemistries to develop a therapeutically useful compound. One approach is to develop a biomarker for toxicity so that systemic exposure can be detected at low levels without encountering severe adverse effects. Secondly, since XEN602 has a very short half-life of about 30 min, it is only necessary to slow systemic uptake sufficiently so that there is little accumulation of drug in the plasma and an extended release formulation might be useful. Thirdly, it may be possible to append much larger substituents onto the active pharmacophore to minimize systemic exposure under a broader range of conditions. These strategies are not mutually exclusive and it may be possible to combine all three to produce a therapeutically useful treatment for iron overload associated with both hereditary hemochromatosis and β-thalassemia intermedia.

## Nonstandard abbreviations

ANOVA: analysis of variance
aPTT: activated partial thromboplastin time
BCECF: 2’,7’-*bis*-(2-carboxyethyl)-5-(And-6)-carboxyfluorescein, a pH indicator dye
CFTR: cystic fibrosis transmembrane conductance regulator
CHO: Chinese hamster ovary
CYP3A4: cytochrome P450 3A4, an enzyme important for drug metabolism
DMT1: divalent metal transporter 1
GI: gastrointestinal
hDMT1: human DMT1
HEK: human embryonic kidney
HH: hereditary hemochromatosis
IV: intravenous
MES: 4-morpholineethanesulfonic acid
Papp: apparent permeability coefficient
p.o.: per os (orally)
PT: prothrombin time
Tet: tetracycline

## ACKNOWLEDGMENTS

These studies were supported in part by a research grant from Genome British Columbia. At the time of their contribution, all authors were employees of Xenon Pharmaceuticals, Burnaby, BC, Canada. The authors acknowledge Nina Hamilton for editorial support.

## SUPPLEMENTARY DATA

### Supplementary Tables

**TABLE S1.**
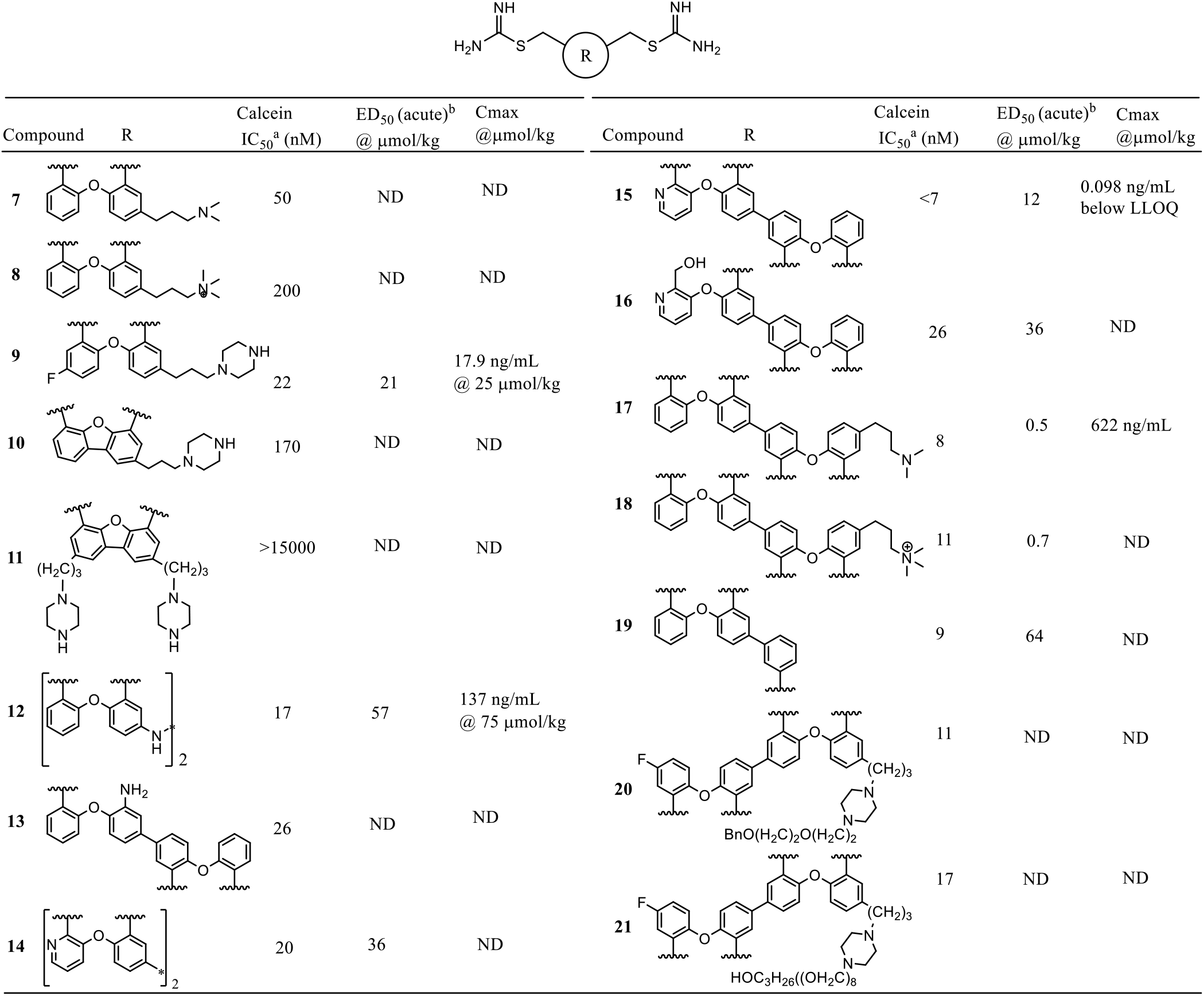
Potency and permeability of DMT1 inhibitors In vitro DMT1 inhibition data and in vivo ED_50_ of iron absorption in an acute model. LLOQ, lower limit of quantitation; ND, not detected.

**TABLE S2.** In vitro pharmacological profiling of **XEN602** and **XEN601** Results for assays with >50% inhibition of specific binding at 1 μM. Drug affinity was determined using standard radioligand binding assays by CEREP SA (Celle L’Evescault, France). α1, XXX; D_1_, XXX; Ca_V_2.2, XXX; DAT, XXX; SERT, XXX. A dash indicates XXX.

| Assay | XEN602<br>% inhibition at 1 μM | XEN601<br>% inhibition at 1 μM |
| --- | --- | --- |
| $\alpha_1$ (non-selective) | 44 | 51 |
| D<sub>1</sub> | 59 | 69 |
| Muscarinic (non-selective) | 56 | -- |
| Ca<sub>v</sub>2.2 | 97 | 82 |
| DAT | 66 | 73 |
| SERT | 85 | 70 |
| choline transporter | 52 | 62 |

### Supplementary Figure

**Fig. S1.**
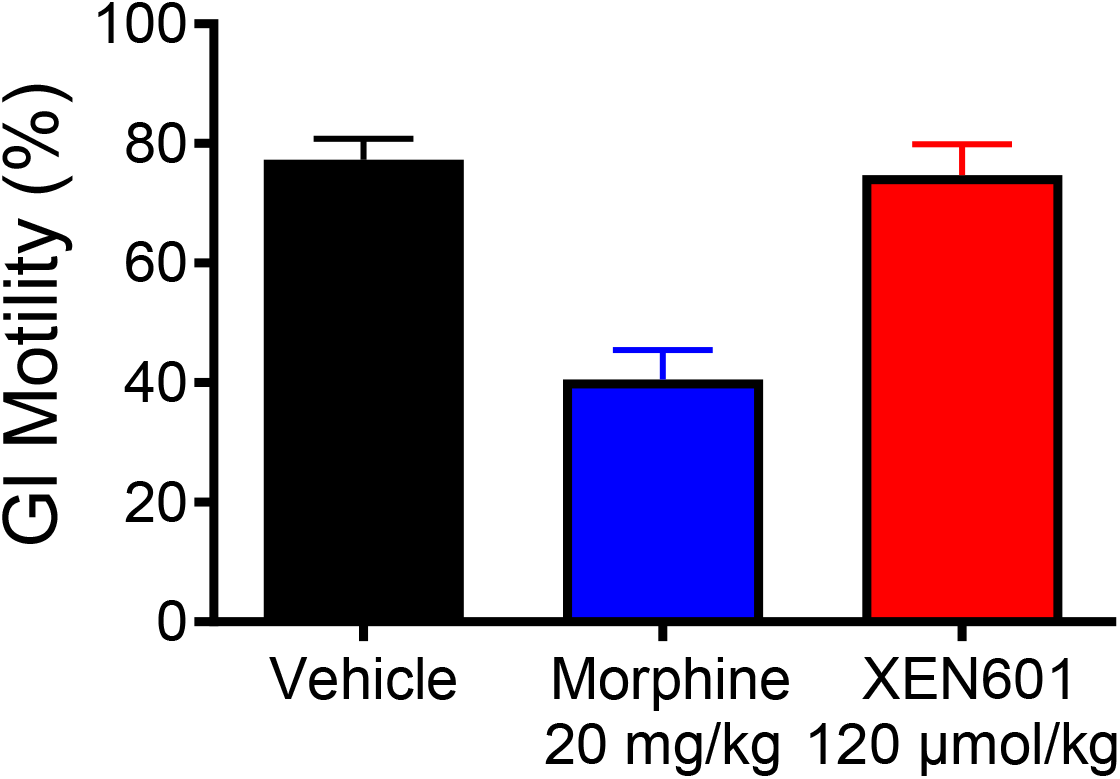
**XEN601** does not influence gut transit time in a rodent model. Intestinal transit of charcoal was measured in Sprague Dawley rats at Ricerca Biosciences (Peitou, Taiwan) using their standard protocol. The test groups consisted of vehicle, 20 mg/kg morphine (a positive control) and 100 mg/kg **XEN601**.

## REFERENCES

Andrews NC and Schmidt PJ (2007) Iron homeostasis. Annu Rev Physiol 69:69–85.

Artursson P, Palm K, and Luthman K (2001) Caco-2 monolayers in experimental and theoretical predictions of drug transport. Adv Drug Deliv Rev 46:27–43.

Breuer W, Epsztejn S, Millgram P, and Cabantchik IZ (1995) Transport of iron and other transition metals into cells as revealed by a fluorescent probe. Am J Physiol 268:C1354–1361.

Brown JX, Buckett PD, and Wessling-Resnick M (2004) Identification of small molecule inhibitors that distinguish between non-transferrin bound iron uptake and transferrin-mediated iron transport. Chem Biol 11:407–416.

Buckett, PD and Wessling-Resnick M (2009) Small molecule inhibitors of divalent metal transporter-1. Am J Physiol Gastrointest Liver Physiol 296:G798–804.

Cadieux JA, Zhang Z, Mattice M, Brownlie-Cutts A, Fu J, Ratkay LG, Kwan R, Thompson J, Sanghara J, Zhong J, et al. (2012) Synthesis and biological evaluation of substituted pyrazoles as blockers of divalent metal transporter 1 (DMT1). Bioorg Med Chem Lett 22:90–95.

Cellier MFM (2012) Nramp: from sequence to structure and mechanism of divalent metal import. Curr Top Membr 69:249–293.

Ehrnstorfer IA, Geertsma ER, Pardon E, Steyaert J, and Dutzler R. (2014) Crystal structure of a SLC11 (NRAMP) transporter reveals the basis for transition-metal ion transport. Nat Struct Mol Biol 21:990–996.

Fleming MD, Romano MA, Su MA, Garrick LM, Garrick MD, and Andrews NC (1998) *Nramp2* is mutated in the anemic Belgrade (*b*) rat: evidence of a role for *Nramp2* in endosomal iron transport. Proc Natl Acad Sci U S A 95:1148–1153.

Fleming MD, Trenor CC 3rd, Su MA, Foernzler D, Beier DR, Dietrich WF, and Andrews NC (1997) Microcytic anaemia mice have a mutation in Nramp2, a candidate iron transporter gene. Nat Genet 16:383–386.

Fleming RE and Ponka P (2012) Iron overload in human disease. N Engl J Med 366:348–359.

Gardenghi S, Ramos P, Marongiu MF, Melchiori L, Breda L, Guy E, Muirhead K, Rao N, Roy CN, Andrews NC, et al. (2010) Hepcidin as a therapeutic tool to limit iron overload and improve anemia in β-thalassemic mice. J Clin Invest 120:4466–4477.

Garrick MD and Garrick LM (2009) Cellular iron transport. Biochim Biophys Acta 1790:309–325.

Gunshin H, Fujiwara Y, Custodio AO, Direnzo C, Robine S, and Andrews NC (2005) Slc11a2 is required for intestinal iron absorption and erythropoiesis but dispensable in placenta and liver. J Clin Invest 115:1258–1266.

Gunshin H, Mackenzie B, Berger UV, Gunshin Y, Romero MF, Boron WF, Nussberger S, Gollan JL, and Hediger MA (1997) Cloning and characterization of a mammalian proton-coupled metal-ion transporter. Nature 388:482–488.

Kelleher T, Ryan E, Barrett S, Sweeney M, Byrnes V, O’Keane C, and Crowe J (2004) Increased DMT1 but not IREG1 or HFE mRNA following iron depletion therapy in hereditary haemochromatosis. Gut 53:1174–1179.

Lynch SR, Skikne BS, and Cook JD (1989) Food iron absorption in idiopathic hemochromatosis. Blood 74:2187–2193.

Mackenzie B, Ujwal ML, Chang M-H, Romero MF, and Hediger MA (2006) Divalent metal-ion transporter DMT1 mediates both H^+^-coupled Fe^2+^ transport and uncoupled fluxes. Pflugers Arch 451:544–558.

Picard V, Govoni G, Jabado N, and Gros P (2000) Nramp 2 (DCT1/DMT1) expressed at the plasma membrane transports iron and other divalent cations into a calcein-accessible cytoplasmic pool. J Biol Chem 275:35738–35745.

Priwitzerova M, Nie G, Sheftel AD, Pospisilova D, Divoky V, and Ponka, P (2005) Functional consequences of the human DMT1 (SLC11A2) mutation on protein expression and iron uptake. Blood 106:3985–3987.

Shayeghi M, Latunde-Dada GO, Oakhill JS, Laftah AH, Takeuchi K, Halliday N, Khan Y, Warley A, McCann FE, Hider RC, et al. (2005) Identification of an intestinal heme transporter. Cell 122:789–801.

Sonawane ND, Zhao D, Zegarra-Moran O, Galietta LJV, and Verkman AS (2008) Nanomolar CFTR inhibition by pore-occluding divalent polyethylene glycol-malonic acid hydrazides. Chem Biol 15:718–728.

Taher A, Hershko C, and Cappellini MD (2009) Iron overload in thalassaemia intermedia: reassessment of iron chelation strategies. Br J Haematol 147:634–640.

Tenopoulou M, Kurz T, Doulias P-T, Galaris D, and Brunk UT (2007) Does the calcein-AM method assay the total cellular “labile iron pool” or only a fraction of it? Biochem J 403:261–266.

Thomas F, Serratrice G, Béguin C, Aman ES, Pierre JL, Fontecave M, and Laulhère JP (1999) Calcein as a fluorescent probe for ferric iron. Application to iron nutrition in plant cells. J Biol Chem 274:13375–13383.

Wetli HA, Buckett PD, and Wessling-Resnick M (2006) Small-molecule screening identifies the selanazal drug ebselen as a potent inhibitor of DMT1-mediated iron uptake. Chem Biol 13:965–972.

Xu H, Jin J, DeFelice LJ, Andrews NC, and Clapham DE (2004) A spontaneous, recurrent mutation in divalent metal transporter-1 exposes a calcium entry pathway. PLoS Biol 2:e50.

Zhang A-S, Canonne-Hergaux F, Gruenheid S, Gros P, and Ponka P (2008) Use of *Nramp2*-transfected Chinese hamster ovary cells and reticulocytes from *mk/mk* mice to study iron transport mechanisms. Exp Hematol 36:1227–1235.

Zhang Z, Kodumuru V, Sviridov S, Liu S, Chafeev M, Chowdhury S, Chakka N, Sun J, Gauthier SJ, Mattice M, et al. (2012) Discovery of benzylisothioureas as potent divalent metal transporter 1 (DMT1) inhibitors. Bioorg Med Chem Lett 22:5108–5113.

